# The Universal Receptive System acts as a novel regulator in the production of antimicrobial and anticancer bioactive compounds by white blood cells

**DOI:** 10.1101/2024.12.17.628964

**Authors:** Victor Tetz, Kristina Kardava, Maria Vecherkovskaya, Alireza Khodadadi-Jamayran, Aristotelis Tsirigos, George Tetz

## Abstract

Despite recent advances, the regulation of anticancer and antimicrobial bioactive compound (AABC) production by leukocytes remains poorly understood. Here, we demonstrate that inactivation of the DNA- and RNA-based Teazeled receptors of the Universal Receptive System in human leukocytes generated so called “Leukocyte-Tells,” which showed enhanced AABC production. Comprehensive analysis of the AABCs produced by Leukocyte-Tells based on LC/MS identified 707 unique or differentially produced peptide or non-peptide metabolites. Functional testing demonstrated that many of these metabolites exhibited increased antibacterial, antifungal, and anticancer activities.

The AABCs produced by the Leukocyte-Tells were effective against different multidrug-resistant clinical isolates of fungi and gram-positive and gram-negative bacteria (including their biofilms), as well as various cancer cell lines, with >100,000-fold activity than AABCs derived from control leukocytes. Notably, the AABCs produced by the Leukocyte-Tells exhibited greater stability under adverse environmental conditions. Our findings highlight the important role of the Universal Receptive System in regulating AABC production through a process named here as cell genome memory management, offering new insights into immune functions and suggesting potential therapeutic applications.

**Summary:** The Universal Receptive System acts as a novel regulator of biosynthetic activity in leukocytes. Modulating the leukocyte Universal Receptive System by inactivating Teazeled receptors triggers the production of new compounds not observed in naïve cells. We refer to these TezR-modified cells as “Leukocyte-Tells.” Leukocytes produce unique metabolites with strong anticancer and antimicrobial activities. Reproducibility in generating leukocytes from the blood of different donors suggests that the observed alterations in cell activity were preprogrammed.

## Introduction

Leukocytes, including granulocytes and agranulocytes, are key components of the innate immune system that can detect and eliminate pathogens and abnormal cells (1). The production of anticancer and antimicrobial products constitutes a fundamental defense mechanism that contributes to immune responses against various bacterial, viral, fungal, and parasitic infections, as well as cancer cells (2–4).

Anticancer and antimicrobial bioactive compounds (AABCs) can be divided into several categories, such as peptides, proteins, and metabolites, based on their chemical structures (5).

Despite modern techniques, such as high-throughput screening, bioinformatics analyses, and advanced proteomic technologies, only a small number of AABCs from human leukocytes have been extensively studied, and the functions of many other products remain largely unknown (1). Traditional research on human leukocyte-derived AABCs has primarily focused on chemokines, antibodies, cytokines, and molecules with direct anticancer and antimicrobial activities, such as myeloperoxidases, lysozymes, integrins, and defensins (known for their antimicrobial activities against various pathogens and their immunomodulatory effects) (5–10). However, the roles of many other factors, particularly those that indirectly antagonize cancer and microorganisms, have not been sufficiently studied (11).

Although AABCs undeniably serve important roles in defending against infections and cancers, they often do not provide adequate protection (12). One reason for this lack of protection is that microbial strategies can reduce the efficacy of AABCs with direct cytotoxic mechanisms. These strategies include modifications to membrane and cell wall structures, alterations in transport systems, and the use of efflux pumps, which help microorganisms evade the cytotoxicity of antimicrobial peptides (13, 14).

Furthermore, the regulation of AABC production by leukocytes, which involves a complex interplay between various signaling pathways, transcription factors, and cellular processes, also influences the effectiveness of protection provided by AABCs (14). Factors such as microbial exposure, inflammatory signals, and the tissue microenvironment can influence the expression and secretion of AABCs by leukocytes (12, 15). Recent findings have highlighted the roles of epigenetic mechanisms, post-transcriptional modifications, and intercellular communication networks in modulating AABC production in response to various stimuli (10, 16). Despite their importance and advances in our understanding of their underlying biology, the exact mechanisms regulating leukocyte AABC production remain unclear.

Until recently, traditional biological approaches for directing cell-based production of desired products were limited, mainly relying on modifications to cellular environments and genome-editing techniques (17–19). These approaches have been used in various contexts ranging from natural biological systems to industrial biomanufacturing.

We recently discovered two previously unknown functions of DNA and RNA as part of the “New Biology” that are distinct from their conventional roles in encoding genetic information and synthesizing proteins. These novel functions involve DNA and RNA acting as a part of the extrabiome (20). One of these functions, termed as the “Pliers function,” involves direct interactions between nucleic acids and proteins that result in the formation of novel protein isoforms including those with prionogenic properties (21–23). We also found that DNA and RNA located outside the cell membrane form DNA- and RNA-based Teazeled receptors (TezRs), which exhibit receptive and regulatory functions (24, 25). The involvement of reverse transcriptases and recombinases in signaling occurring downstream of TezRs constitutes a Universal Receptive System (24, 25). Additionally, the receptive role of DNA was assessed in artificially constructed DNA receptors (26).

Subsequently, we demonstrated that the Universal Receptive System can orchestrate various cellular behaviors and facilitate the emergence of novel properties across a wide range of prokaryotic and eukaryotic cells. Moreover, this system plays key regulatory roles not only at the single-cell level but also in multicellular processes, such as regulating plant growth (24). Using various cellular model systems, we highlighted the effectiveness of manipulating the Universal Receptive System in orchestrating key cellular features. Specifically, we showed the essential role of this system in regulating cellular responses to a broad range of chemical, physical, and biological stimuli, as well as in controlling cell memory and forgetting (25, 27). Additionally, we showed that the loss of DNA- and RNA-based TezR altered the secretion profiles of various bacterial and fungal enzymes, including those essential for breaking down nutrient substrates (24).

In this study, we demonstrate that modulating the Universal Receptive System through DNA- and RNA-based TezRs results in the acquisition of enhanced properties by human leukocytes. Here, we introduce the term “Leukocyte-Tells” to refer to these modified cells. Notably, our results revealed that these modified immune cells exhibited enhanced production of AABCs with unique compositions and properties. Our study represents an initial assessment of how cell genome-memory management enables the Universal Receptive System to orchestrate AABC production in leukocytes, revealing novel regulatory mechanisms underlying immune functions.

## Results

### AABC production by Leukocyte-Tells

First, we investigated the differences in secretion of proteins implicated in immune cells activation and those having direct antimicrobial or anticancer activities by Leukocyte-Tells in response to LPS stimulation. We identified 8 proteins that were either uniquely identified or exhibited statistically significant increases in relative abundance (p < 0.05) in the culture media of Leukocyte-Tells compared to Leukocyte-Controls(Table 1).

**Table 1.** Enriched and unique proteins in supernatants of Leukocyte-Tells associated with immune cell activation, antimicrobial, and anticancer activity.

| Protein ID | Protein name | Log2 fold change | Functions |
| --- | --- | --- | --- |
| **Immune cell activation, antimicrobial and anticancer activity** |  |  |  |
| AZU1 | Azurocidin | 2.263 | Antimicrobial & Inflammation Modulation |
| CAT | Catalase | 0.876 | Hydrogen Peroxide Signaling |
| CLC | Galectin-10* | 3.459 | Regulates T Cell & Dendritic Cell Functions |
| LCN2 | Neutrophil gelatinase-associated lipocalin | 3.776 | Sequesters Iron to Inhibit Bacterial Growth |
| LYZ | Lysozyme | 1.678 | Antimicrobial activity, cell activation |
| MPO | Myeloperoxidase | 1.7 | Antimicrobial, immune activation, anticancer activity |
| S100A12 | Protein S100-A12 | 4.087 | Direct antimicrobial binding, disrupting cell integrity |
| S100A9 | Protein S100-A9 | 0.907 | Acts as DAMP signaling inflammation |

\* Uniquely found in leukocyte supernatants

Within this group, many proteins associated with immune cell activation and antimicrobial and anticancer activities were observed, such as azurocidin (AZU1), lysozyme (LYZ), myeloperoxidase (MPO), and protein S100-A12 (S100A12) and others (28–32). These proteins have direct antimicrobial and anticancer functions, highlighting their roles as hallmarks of immune cells actively responding to bacterial infections or inflammation (33, 34).

Next, we compared the metabolomes of AABCs produced by Leukocyte-Tells and Leukocyte-Controls.

Of the 496 differentially produced metabolites identified using untargeted metabolomics methods, 285 were short peptides (180 unique and 185 present at significantly higher abundances) (Fig. 1A, Table S1).

**Figure 1.**
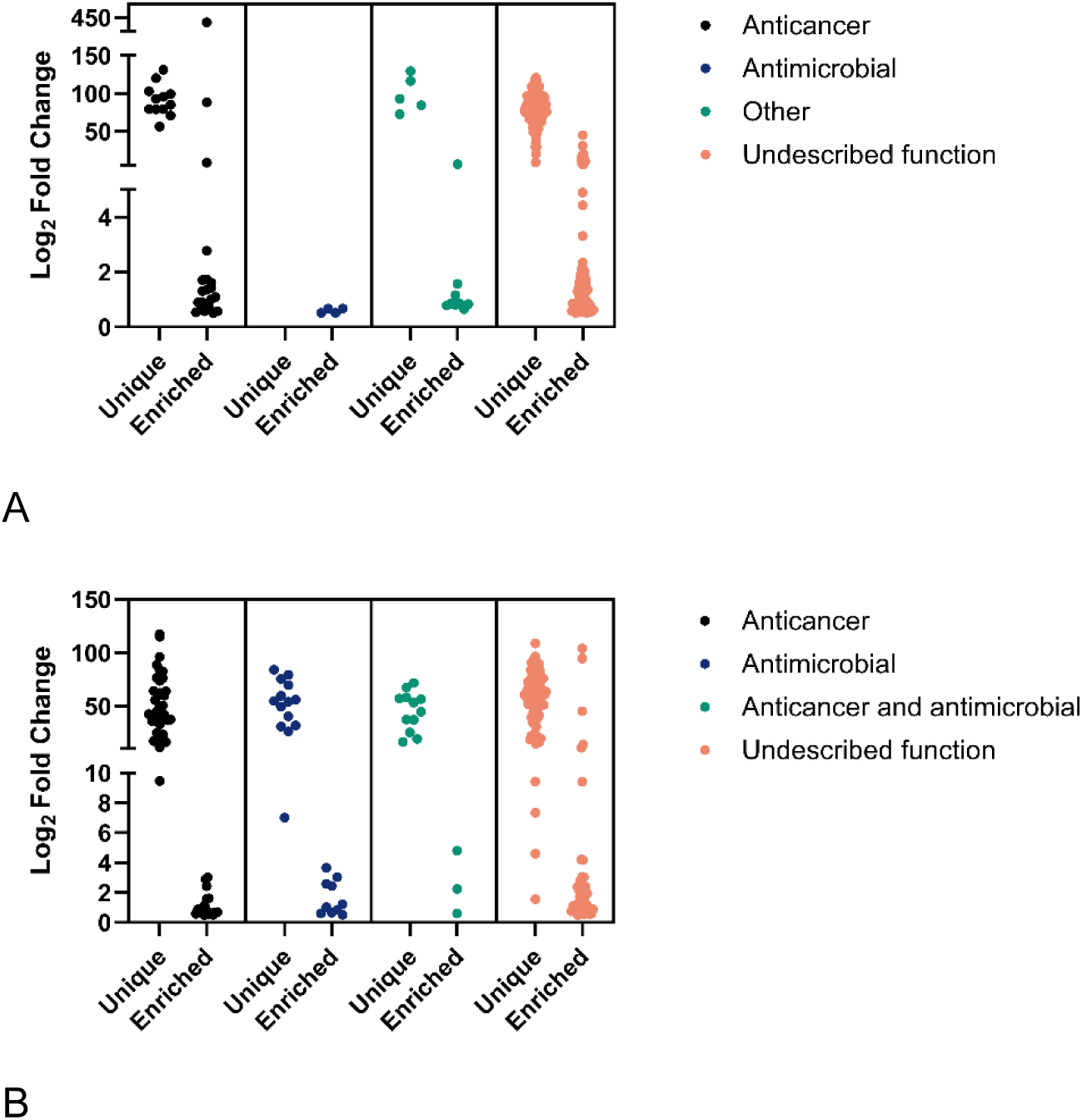
Enriched and unique metabolites in the supernatants of Leukocyte-Tells. These metabolites included (A) short peptides and (B) non-peptide metabolites.

Of these metabolites, 32 have known anticancer properties. Some of them are angiotensin I-converting enzyme inhibitor (e.g., Ala–Ile, Asp–Met, His–Phe, Ser–Tyr, Val– Trp, and others), which have been shown to slow tumor progression and inhibit tumor-associated angiogenesis and metastasis (35, 36). Diffusible bitter peptides (Arg–Leu, Gly–Leu, Leu–Lys, Pro–Val, and Val–Phe) promote apoptosis in gastrointestinal and head and neck cancers (37). Six peptides (Ile–Arg, Lys–Ala, Lys–Lys, Pro–Ala, Pro–Pro– Gly, and Val–Val) match bioactive peptides from amaranth seeds, which can suppress cell-migration capabilities and induce apoptosis in breast cancer (38, 39). Two peptides (Glu–Tyr and Trp–Phe), matching the cyano-phycocyanin (C-PC) protein, were extracted from *Oscillatoria minima* and shown to have antimicrobial, algicidal, and antiradical activities (40). The 14 oligopeptides have various effects, including neuroactive (Phe– Tyr), hepatoprotective (Leu–Glu), and antihypertensive effects (Phe–Gln). The remaining peptides have no known biological functions, and their roles as leukocyte metabolites remain unclear.

A group of 211 non-peptide metabolites, uniquely found in the supernatants of Leukocyte-Tells (141 metabolites) and among differentially expressed metabolites, comprised complex molecules with various chemical properties (Fig. 1B, Table S2).

Among these metabolites, 54 have been shown to exhibit anticancer activity. These compounds, which predominantly comprise alkaloids, heterocyclic compounds, phenolic compounds, and aromatic compounds, can target various cellular functions through mechanisms such as enzyme inhibition (e.g.,1-benzylimidazole), interference with mutant peptides (e.g., 1,2-di(2-pyridyl)ethylene), inducing apoptosis (e.g., lumichrome), or toxicity against fast-dividing cancer cells (e.g., desacetylvindoline and vincristine) (41–44). In addition, 24 of the metabolites have antimicrobial activity against bacteria (leoidin dimethyl ether) and protists (2-[1-[(4-chlorophenyl)-oxomethyl]-5-methoxy-2-methyl-3-indolyl]-1-(4-morpholinyl)ethanone), and some metabolites are leukotrienes that regulate the immune response (such as N-acetyl-LTE4) (45–47).

Furthermore, a subset of 15 metabolites was identified that possesses both anticancer and antimicrobial activities, mostly due to ATP depletion (α-jasmonic acid), apoptosis induction (lauric acid), or targeting mitochondrial metabolism (5-ethyl-2′-deoxyuridine) (48–52). Another metabolite (2-(4-isobutylphenyl)propanoic acid) has non-steroidal anti-inflammatory drug activity and limit cell migration and increase chemo-sensitivity in the tumor microenvironment (53).

Most other chemicals have various biological functions, including neuroprotection (dexanabinol), and include vitamins, amino acids, and prostaglandins (54).

### AABCs derived from Leukocyte-Tells had superior antimicrobial effects

First, we assessed the antimicrobial activity of AABCs derived from various leukocyte preparations. Supernatants from 24 h cultures of Leukocyte-Control and Leukocyte-Tells were evaluated using the disk diffusion method (55). Paper disks immersed in AABC-containing supernatants were air-dried and placed on Petri dishes containing the test microbes. The results summarized in Fig. 2A show the mean values of the growth-inhibition zones. Notably, AABCs derived from Leukocyte-Tells consistently exhibited significantly larger inhibition zones than those derived from Leukocyte-Controls across all tested strains, including antibiotic-resistant clinical isolates, indicative of their superior antimicrobial activities (Fig. 2A, B).

**Figure 2.**
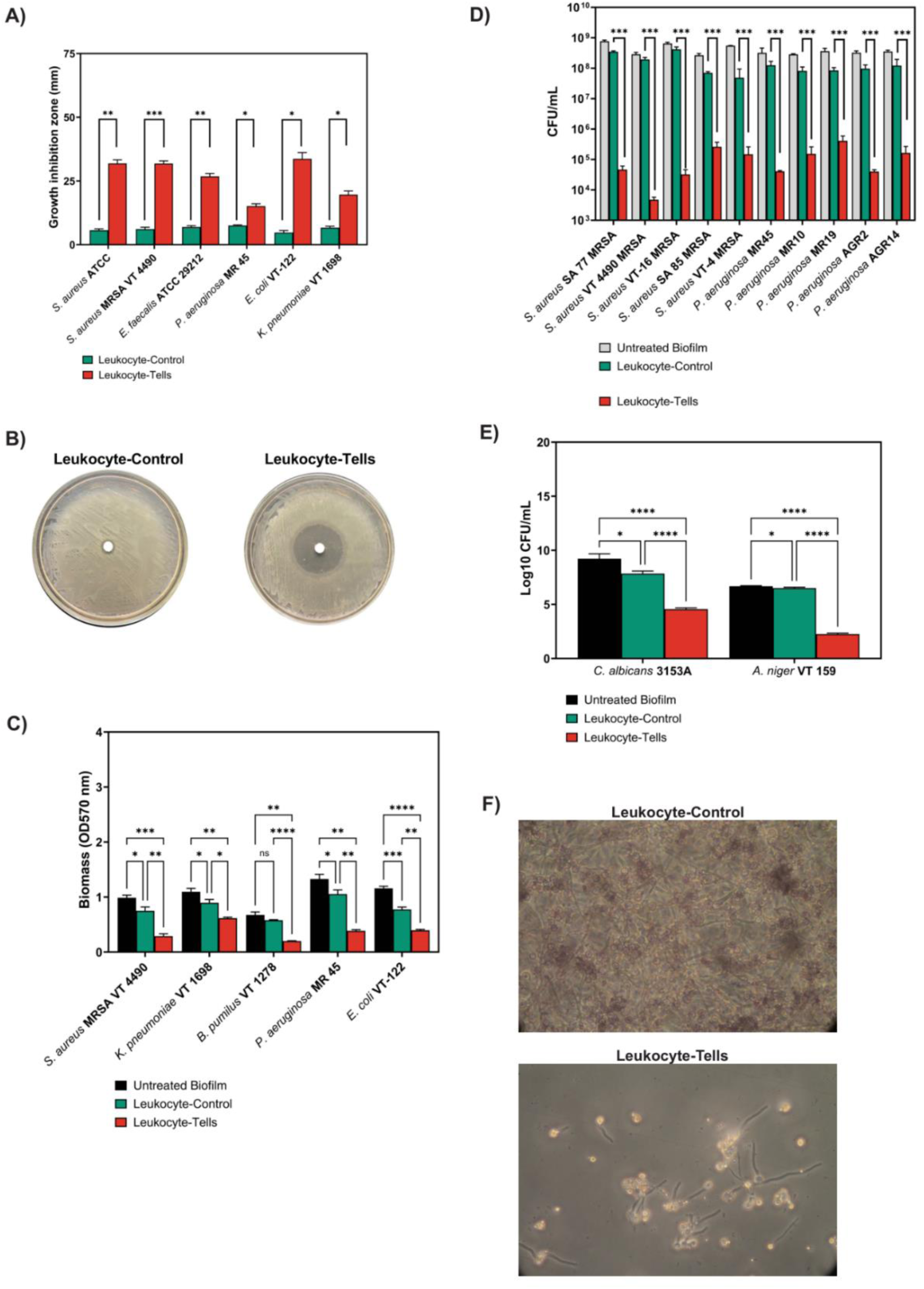
Antimicrobial activity of AABCs derived from Leukocyte-Tells. (A) Growth-inhibition zones observed with various microbial cultures after treatment with AABCs derived from Leukocyte-Controls and Leukocyte-Tells using a disk diffusion method. The combined results of three independent experiments are depicted. The bar graphs represent the mean ± standard error of the mean (SEM). Statistical significance was determined via two-way analysis of variance (ANOVA). (B) Examples of growth-inhibition zones observed after treating *E. coli* VT-122 cells with supernatants containing AABCs - from Leukocyte-Control and Leukocyte-Tell cultures. (C) Inhibition of 24 h old bacterial biofilms with undiluted supernatants containing AABCs derived from Leukocyte-Control and Leukocyte-Tell cultures. The combined results of three independent experiments are depicted. The bar graphs represent the mean ± SEM. Statistical significance was determined via two-way ANOVA. (D) Decreased CFUs of 24 h old bacterial biofilms treated with supernatants containing AABCs derived from Leukocyte-Controls and Leukocyte-Tells. The combined results of three independent experiments are depicted. The bar graphs represent the mean ± SEM. (E) Decreased CFUs of 24 h old fungal biofilms treated with supernatants containing AABCs derived from Leukocyte-Controls and Leukocyte-Tells. The combined results of three independent experiments are depicted. The bar graphs represent the mean ± SEM. Statistical significance was determined via two-way ANOVA. (F) Changes in the morphology and density of *A. niger* VT 159 biofilms after treatment with supernatants containing AABCs derived from Leukocyte-Control and Leukocyte-Tell cultures.

We further evaluated the efficacies of AABCs derived from various leukocyte sources based on the minimal inhibitory concentration (MIC), where the MIC represents the highest dilution (the minimal concentration) of an AABC required to completely inhibit microbial growth (56). To ensure that the observed antimicrobial effects were a direct consequence of DNA- and RNA-based TezR inactivation, we introduced an additional control in which TezRs were neutralized with anti-DNA and anti-RNA antibodies instead of being destroyed by nucleases. In these settings, AABCs derived from Leukocyte-Tells showed greater activity against all tested bacterial and fungal strains than the controls (Table 2).

**Table 2.** MICs of the AABC-containing supernatants.

| Tested strain | Maximal dilution of AABC to reach MIC |  |  |  |
| --- | --- | --- | --- | --- |
|  | Leukocyte-Control | Leukocyte-Tells | Leukocyte-Tells-Ab | IgG Isotype Control |
| *S. aureus* ATCC 29213 | 1:2 | 1:4 | 1:4 | n.a. |
| *S. aureus* VT 209R | n.a. | 1:2 | 1:2 | n.a. |
| *S. aureus* MRSA VT 4490 | n.a. | 1:2 | 1:2 | n.a. |
| *B. pumilus* VT 1278 | n.a. | 1:2 | 1:2 | n.a. |
| *E. faecalis* ATCC 29212 | n.a. | 1:2 | 1:2 | n.a. |
| *P. aeruginosa* MR45 | n.a. | 1:2 | -* | n.a. |
| *E. coli* VT 122 | n.a. | 1:4 | -* | n.a. |
| *K. pneumoniae* VT 1698 | n.a. | n.a. | -* | n.a. |
| *C. albicans* 3153A | n.a. | 1:4 | -* | n.a. |
| *A. niger* VT 1591 | n.a. | 1:4 | -* | n.a. |

Bacterial and fungal strains were treated with AABC-containing supernatants derived from Leukocyte-Control and Leukocyte-Tell cultures. The Leukocyte-Tell-Ab control was generated by treating leukocytes with anti-RNA and anti-DNA antibodies. IgG Isotype was used as a control for Leukocyte-Tell-Ab. The data presented are from three independent experiments. \*not performed. n.a., not applicable; the undiluted supernatant that did not achieve the MIC.

Undiluted AABC samples derived from Leukocyte-Control cells did not reach the MIC or completely inhibit the growth of any microorganisms tested, except for *S. aureus* ATCC 29213, whereas those derived from Leukocyte-Tells achieved the MIC within a 1:2 to 1:4 dilution range against all bacteria and fungi, except for *K. pneumoniae*. Importantly, no difference was found in the efficacy of AABCs derived from leukocytes treated with nucleases (Leukocyte-Tells) and those treated with anti-DNA and anti-RNA antibodies, strongly suggesting that the observed antimicrobial activity was a direct consequence of TezR system disruption (Table 2). As expected, the IgG isotype control showed no detectable antimicrobial activity.

Next, we assessed the effects of AABCs on preformed biofilms, which frequently exhibit enhanced resistance to immune responses (57, 58). Treatment with undiluted AABCs from Leukocyte-Tells led to a 50–70% decrease in the biofilm biomass and a 10,000–100,000-fold decrease in viable counts relative to intact biofilms (Fig. 2C, D, Fig. S1A, B). This pattern was observed in all 24 and 48 h old biofilms derived from gram-positive and gram-negative bacteria. Importantly, a similar trend was observed across all tested strains, including 10 multidrug-resistant clinical isolates of *S. aureus* and *P. aeruginosa* isolated from patients with severe lung infections. In contrast, treatment with AABCs derived from Leukocyte-Control cells only induced a minimal reduction in bacterial growth and an insignificant decrease in viable cell counts.

Similarly, treatment with AABCs derived from Leukocyte-Tells substantially inhibited fungal biofilm formation with both *C. albicans* 3153A and *A. niger* VT 1591, resulting in a 1,000–100,000-fold lower colony-forming units (CFUs) than controls. In contrast, AABCs derived from Leukocyte-Controls showed minimal to no fungal biofilm inhibition (Fig. 2E, Fig. S1B). Post-treatment microscopic evaluation of *A. niger* biofilms reinforced these findings, clearly demonstrating a strong reduction in biofilm formation after treatment with AABCs from Leukocyte-Tells (Fig. 2F).

### In vitro anticancer activity of AABCs derived from Leukocyte-Tells

To evaluate the anticancer activities of AABCs derived from various leukocyte sources, we incubated a human non-small cell lung cancer (NSCLC) cell line (NCI-H1299 cells) and a glioblastoma cell line (T98G cells) with serial dilutions of supernatants from 24 h cultures of Leukocyte-Control and Leukocyte-Tells. The metabolic properties of cancer cells were evaluated by performing standard 3-(4,5-dimethylthiazol-2-yl)-2,5-diphenyltetrazolium bromide (MTT) assays, and cancer cell viability was measured using a colorimetric lactate dehydrogenase (LDH) assay in the presence of different concentrations of AABCs.

Notably, supernatants derived from Leukocyte-Controls only had a marginal effect on the metabolic properties (Fig. 3A) and viability (Fig. 3B) of NCI-H1299 cells. In stark contrast, even 100-fold diluted AABC supernatants derived from Leukocyte-Tells induced >50% metabolic inhibition and decreased the viability of the NSCLC cells (Fig. 3A, B).

**Figure 3.**
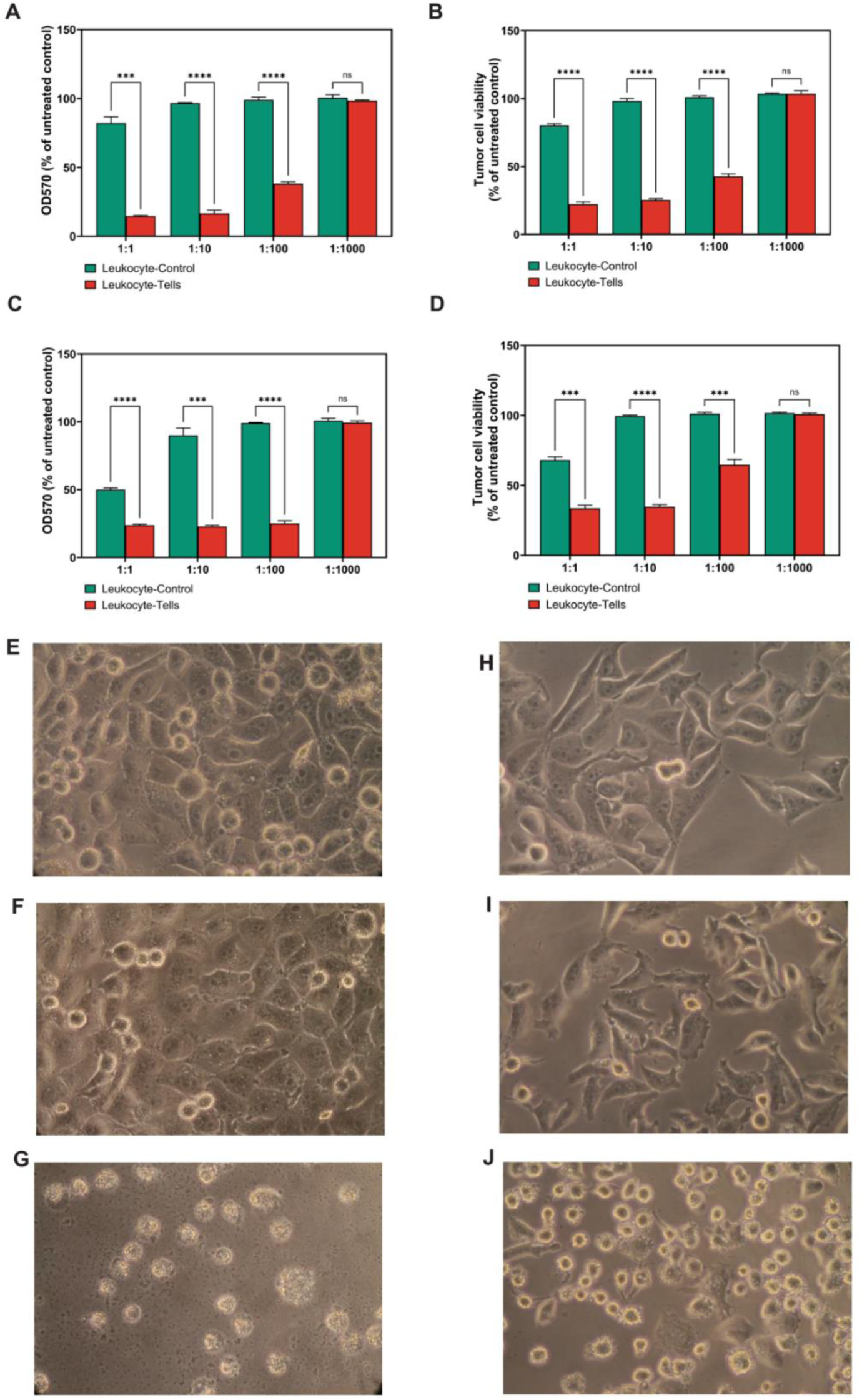
Anticancer activity of Leukocyte-Tell-derived AABCs. (A–D) Metabolic properties and viability of the NCI-H1299 and T98G cancer cell lines incubated with various dilutions of supernatants containing AABCs derived from Leukocyte-Control and Leukocyte-Tell cultures. The metabolic properties and viability of NCI-H1299 and T98G cells were assessed by performing standard MTT (A, C) and colorimetric LDH (B and D) assays, respectively. The combined results of three independent experiments are depicted. The bar graphs represent the mean ± SEM. Statistical significance was determined by performing unpaired t-tests. (E, H) Growth of untreated NCI-H1299 and T98G cells. (F, I) Growth of NCI-H1299 and T98G cells after treatment with supernatants containing AABCs derived from Leukocyte-Controls. (G, J) Growth of NCI-H1299 and T98G cells after treatment with supernatants containing AABCs derived from Leukocyte-Tells.

Although T98G cells appeared more sensitive to supernatants from Leukocyte-Controls than NCI-H1299 cells (as indicated by a 50% reduction in metabolic activity and a 25% reduction in viability), this effect was only observed with undiluted supernatants (Fig. 3C, D). Conversely, even a 100-fold dilution of Leukocyte-Tell-derived supernatant decreased the metabolic activity of glioblastoma cells by 75% and decreased their viability by 25%.

These findings were further strengthened by post-treatment microscopy assessments, which showed clear disruptions in monolayers of both the NSCLC and glioblastoma cell lines in the presence of Leukocyte-Tell-derived supernatants, whereas mostly intact monolayers were observed in the presence of Leukocyte-Control-derived supernatants (Fig. 3E, F).

### AABCs produced by Leukocyte-Tells exhibited unique temperature and protease resistance

Previously, we demonstrated that the Universal Receptive System regulated cellular responses to high temperatures and proteolytic digestion in both prokaryotic and eukaryotic systems (25). Based on these findings, we assessed the resistance of AABCs derived from various leukocyte sources to environmental stressors. Specifically, various dilutions of supernatants containing AABCs exposed to elevated temperatures (70°C or 90°C) or pe-treated with proteolytic enzymes (trypsin and proteinase K) were added to *S. aureus* ATCC 29213 cells as the only bacterial strain sensitive to AABCs from Leukocyte-Controls. The maximal dilutions required to achieve the MIC were determined.

AABC-containing supernatants derived from Leukocyte-Control lost their antimicrobial properties upon heating. Therefore, after exposure to 90°C for 30 min, even the undiluted samples from the Leukocyte-Controls were unable to achieve the MIC (Table 3). In stark contrast, AABC-containing supernatants from Leukocyte-Tells consistently reached the MIC, even when subjected to multiple-fold dilutions across all evaluated temperatures and exposure times studied (Table 3).

**Table 3.** Maximal dilution of AABC-containing supernatants after heat expose required to reach the MIC.

| AABC source | Maximal dilution of AABC to reach MIC for *S. aureus* ATCC 29213 |  |  |  |  |  |  |
| --- | --- | --- | --- | --- | --- | --- | --- |
|  | RT | 70°C |  |  | 90°C |  |  |
|  |  | 5 min | 15 min | 30 min | 5 min | 15 min | 30 min |
| Leukocyte-Control | 1:2 | 1:2 | 1:2 | 1:2 | 1:2 | 1:2 | >1:1 |
| Leukocyte-Tells | 1:8 | 1:8 | 1:8 | 1:8 | 1:8 | 1:8 | 1:8 |

AABCs derived from Leukocyte-Controls were sensitive to treatment with trypsin or proteinase K, which resulted in the complete loss of their antimicrobial activity. Specifically, we detected the remaining activity of AABC-exposed bacteria only after 30 s of incubation with trypsin, and the bacteria completely lost their activity with longer incubation times (Table 4).

**Table 4.** Maximal dilution of AABC-containing supernatants pre-incubated with proteolytic enzymes.

| AABC source | Maximal dilution of AABC to reach MIC for *S. aureus* ATCC 29213 |  |  |  |  |  |  |  |  |
| --- | --- | --- | --- | --- | --- | --- | --- | --- | --- |
|  | PBS | Trypsin |  |  |  | Proteinase K |  |  |  |
|  |  | 0.5 min | 2 min | 15 min | 30 min | 0.5 min | 2 min | 15 min | 30 min |
| Leukocyte-Control | 1:2 | 1:2 | >1:1 | >1:1 | >1:1 | >1:1 | >1:1 | >1:1 | >1:1 |
| Leukocyte-Tells | 1:8 | 1:8 | 1:4 | 1:4 | 1:2 | 1:8 | 1:2 | 1:2 | 1:2 |

This observation was even more pronounced after pretreatment with proteinase K, where no activity was detected regardless of the duration of treatment, even with undiluted supernatants (Table 4). In contrast, AABC-exposed cells derived from Leukocyte-Tells maintained their activity, with only a 2–4 fold reduction in activity, even after prolonged pretreatment with either of the proteolytic enzymes (Table 4).

Taken together, these findings illustrate the enhanced stability and robustness of AABCs derived from Leukocyte-Tells under adverse environmental conditions.

## Discussion

Here, we describe for the first time the role of the Universal Receptive System as a novel orchestrator and regulator of cell synthesis. Modulating this system by inactivating DNA- and RNA-based TezR receptors enabled the development of white blood cells with different synthetic activities that their natural counterparts, and we named these cells Leukocyte-Tells. Our study of AABC production by Leukocyte-Tells revealed substantial differences when compared with Leukocyte-Controls, highlighting various AABC characteristics. These characteristics included differences in the types and quantities of proteins, peptides, and metabolites produced by Leukocyte-Tells, as well as their efficacy against microorganisms and cancer cells.

We identified various supernatant proteins with antimicrobial and anticancer activities that were uniquely identified or whose production was significantly increased in Leukocyte-Tells

The same trend was identified in the metabolomic profiles of supernatants from Leukocyte-Tells, with many of the significantly more abundant metabolites known for their antimicrobial and anticancer activities.

Simultaneously, we identified unusual peptides and metabolites, such as those related to bitter lactones or alkaloids, including glaucarubin and vincristine, which are typically produced by plant cells but not human cells (59, 60).

All untargeted metabolomic identifications were based entirely on MS library database searches against the METLIN database using m/z, MS/MS fragmentation patterns, and isotope similarity scores. The METLIN database is known to include identified metabolites from many species and sample types, not only mammalian cell types. We acknowledge that some of the more exotic identifications have not been confirmed using LCMS analysis of standard compounds, but the fact remains that many of the identified metabolites either specific to or highly upregulated in Leukocyte Tells are known to harbor anti-cancer and anti-microbial function

Collectively, these results suggest that the Universal Receptive System is a novel tool being a part of “New Biology” for activating unusual pathways and altering the cellular capabilities in order to produce novel compounds not observed in naïve cells. We suggest the term “cell-genome memory managing” to emphasize the fact that the induced cell states were pre-programmed and could be reactivated.

Our data confirmed that AABCs from Leukocyte-Tells had a different composition but also demonstrated higher antimicrobial and anticancer activities. In contrast to the AABCs derived from Leukocyte-Controls, which only reach the MIC for *S. aureus*, the AABCs of Leukocyte-Tells exhibited significantly broader antimicrobial activity, being able to inhibit various gram-positive and gram-negative bacteria, including clinical isolates of *S. aureus*, *B. pumilus*, *E. faecalis*, *P. aeruginosa*, and *E. coli*. The only bacterial strain whose growth was not completely inhibited by AABCs from Leukocyte-Tells was the mucoid strain of *K. pneumonia*. These results can be partially explained by the mucoid capsule of *Klebsiella* spp., which may have inactivated AABCs (61). However, in this study, the growth of mucoid *P. aeruginosa* was completely inhibited by AABCs. Importantly, with the disc diffusion method, the AABCs of Leukocyte-Tells significantly inhibited *K. pneumonia* growth. A similar trend was observed for *K. pneumonia* biofilms, where the biomass was significantly inhibited by the AABCs of Leukocyte-Tells.

In the future, we plan to explore why the AABCs showed low activity against *Klebsiella* spp. However, in this study, we were particularly interested in evaluating the antimicrobial activity against biofilms formed by the clinical pathogens, *S. aureus* and *P. aeruginosa*, given their high medical importance (62). Treatment with AABCs from Leukocyte-Tells resulted in up to a 100,000-fold decrease in viable counts (compared with those in intact biofilms) and a 10,000-fold decrease compared with biofilms treated with AABCs derived from Leukocytes-Controls. These results agree with clinical observations that the human immune system frequently cannot overcome biofilms in the body once they are established (57). Additionally, the AABCs produced by Leukocyte-Tells effectively reduced the viable counts in fungal biofilms, which can be even more tolerant to immune cell responses than bacterial biofilms (63–65). Importantly, the efficacy of these AABC remained unaffected by the microorganism’s resistance to antibiotics, as both antibiotic-sensitive and antibiotic-resistant strains (including pan-antibiotic-resistant clinical isolates from patients with cystic fibrosis) responded equally to treatment with AABCs derived from Leukocyte-Tells (66, 67).

The AABCs also exhibited higher activities against various cancer cell lines, including those with mutant p53 or a p53-null profile (68, 69). Taken together, our results suggest that AABC production in Leukocyte-Tells occurs through a p53-independent pathway.

Notably, AABCs derived from Leukocyte-Tells demonstrated greater resilience to proteolytic enzymes, including trypsin and proteinase K, maintaining their biological activity that AABCs isolated from control counterparts. This extended stability underscores the potential of AABCs for robust and sustained antimicrobial and anticancer activities in adverse biological microenvironments (70).

In this study, we used Leukocyte-Tells derived from white blood cells obtained from multiple donors, which revealed a consistent pattern of AABC production across all samples. This uniformity strongly suggests that targeting the Universal Receptive System initiates reproducible changes in cellular characteristics, indicative of a highly regulated and programmed response.

These results illustrate the reliability and predictability of cellular modifications induced by manipulating the Universal Receptive System, highlighting its potential for controlled and standardized applications.

These findings emphasize the potential of the Universal Receptive System for enhancing AABC production and orchestrating the production of various other biologically important molecules across different cell types. The ability to achieve such modulation without genome editing opens opportunities for therapeutic interventions, biotechnological applications, and further research on regulating cellular functions.

## Experimental procedures

### Bacterial and fungal strains and culture conditions

We purchased *Staphylococcus aureus* ATCC 29213 and *Enterococcus faecalis* ATCC 29212 from the American Type Culture Collection (ATCC; Manassas, VA, USA). We also obtained several bacterial and fungal clinical isolates, including *Bacillus pumilus* VT 1278, *Escherichia coli* VT 12, *S. aureus* MRSA VT 4, *S. aureus* MRSA VT 16, *S. aureus* MRSA VT 4490, *S. aureus* MRSA SA 77, *S. aureus* MRSA SA 85, *Klebsiella pneumoniae* VT 1698, *Pseudomonas aeruginosa* MR45, *P. aeruginosa* MR10, *P. aeruginosa* MR19, *P. aeruginosa* AGR2, *P. aeruginosa* AGR 14, *Candida albicans* 3153A, and *Aspergillus niger* VT 159. These isolates were sourced from a private collection provided by the Human Microbiology Institute (New York, NY, USA), samples from patients with cystic fibrosis via the CF Foundation Therapeutics Development Network Resource Center for Microbiology at Seattle Children’s Hospital (Seattle, WA, USA), and Dr. Paul Fidel’s laboratory at Louisiana State University Health Sciences Center (New Orleans, LA, USA).

For experiments involving solid media, bacteria were cultured on Columbia agar; for studies on planktonic-growing bacteria, the bacteria were cultivated in CA-MHB (all materials from Sigma-Aldrich, St. Louis, MO, USA). All cultures were incubated aerobically at 37 °C.

All fungal experiments were conducted using Sabouraud dextrose agar and Sabouraud broth (both from Oxoid, UK), with incubation at 30 °C.

### Cell lines

The H1299 and T98G cell lines (ATCC, Manassas, VA, USA) were authenticated via short tandem repeat profiling. H1299 and T98G cells were cultured in RPMI-1640 medium (Sigma-Aldrich) supplemented with 10% fetal bovine serum (FBS; Gibco, Invitrogen Corporation, NY, USA), 1 mM L-glutamine (Sigma-Aldrich, St. Louis, MO, USA), penicillin G (100 U/mL), and streptomycin (100 mg/mL, Sigma-Aldrich, St. Louis, MO, USA). Cells were cultivated in a humidified chamber (95% air, 5% CO_2_) at 37 °C. Cells were detached by enzymatic treatment by using 0.05% trypsin/EDTA solution (Invitrogen).

### Induction and isolation of AABCs

First, we generated control leukocytes (Leukocyte-Control cells) and Leukocyte-Tells. Leukocytes and platelets were isolated from the buffy coats (BioIVT, PA, USA) of healthy preselected donors who had not taken any antibiotics, corticosteroids, or anticancer drugs for the last 3 months. The leukocytes and platelets were isolates via double gradient centrifugation, as described previously, with Histopaque 1,077 and 1,119 (both from Sigma-Aldrich) per the manufacturer’s instructions (71, 72). The Leukocyte-Control cells comprised untreated leukocytes and platelets. To generate Leukocyte-Tells, the Leukocyte-Control cells were treated with DNase I and RNase A at 1 µg/ml (Thermo Fisher Scientific, USA), as previously described (24).

In some experiments, Leukocyte-Tells were generated by treatment with anti-DNA and anti-RNA antibodies, as previously described (25, 27). Briefly, the supernatants of 24 h *B. pumilus* biofilms were filtered through 0.22 μm filters (Millipore Corp., Bedford, MA, USA) and the extracellular RNA or DNA was extracted using an RNeasy Mini Kit and DNeasy Mini Kit, respectively (Qiagen, Valencia, CA). Antibodies against extracellular RNA or DNA were obtained after immunizing 4-month-old New Zealand white rabbits with nucleic acids and complete Freund’s adjuvant according to an immunization protocol described by Cold Spring Harbor Laboratories (73). To block DNA- or RNA-based receptors generated antibodies were used at 1:3200 dilution. Rabbit IgG Isotype Control (550875, BD, USA) at the same titer was used as control.

Six sets of leukocytes, each sourced from random donors of varying sex, age, and race, were used in this study. All sets were processed without freezing within 24 h of generation.

The leukocyte counts in the samples were calculated using an automated SYSMEX XN-330 hematology analyzer (Sysmex, Japan).

Next, we induced AABC production as previously described with some modifications (74). Briefly, 1 × 10^7^ Leukocyte-Tells or Leukocyte-Control viability >70% were placed in 96-well plates in RPMI-1640 and then incubated with lipopolysaccharide 5 ng/mL; Thermo Fisher) for 3 h at 37 °C. Subsequently, the supernatant was isolated by centrifugation at 150 × *g* for 25 min and passed through a 0.22 µm filter. The resulting supernatant (containing AABCs) was further studied for antimicrobial and anticancer activities on the same day. For the LC/MS and metabolomics studies, the AABCs were frozen at −80 °C until analysis for no longer than 28 days.

### Estimating the MICs for bacteria and the minimum fungicidal concentrations (MFCs) of AABCs in vitro

The MICs for bacteria and MFCs for AABCs were determined using a serial microdilution technique following a modification of the Clinical and Laboratory Standards Institute (CLSI) guidelines (75–77). The standard inocula for bacteria, yeast, and mold cultures were 5 × 10^5^, 2.5 × 10^3^, and 5 × 10^4^ CFUs/mL. We added 20 μL of the bacterial inoculum in CA-MHB broth to separate wells of 96-well plates containing 180 μL of 2-fold serially diluted AABCs in RPMI-1640. The bacterial and fungal cultures were incubated for 24 h at 37 °C or 30 °C, respectively. The MIC and MFC values were defined as the lowest concentration of an AABC that completely inhibited bacterial or fungal growth, respectively. The optical density (OD) of the microorganisms was determined using a GloMax Discover Microplate Reader (Promega, Madison, USA). The MIC and MFC values were normalized to the mean OD_570_ of control wells containing the medium alone.

To assess microbial viability, bacterial and fungal suspensions were serially diluted, and 100 μL of each diluted suspension was spread onto Columbia agar plates (for bacteria) and Sabouraud dextrose agar (for fungi). The plates were incubated overnight at 37 °C (for bacteria) and 30 °C (for fungi), and CFUs were counted the following day. The tests were performed in triplicate in three independent experiments.

### Disk diffusion testing

The susceptibility to AABCs was determined using the Kirby–Bauer disc diffusion method according to criteria defined by the CLSI, with some modifications (55, 78). Sterile disks without antibiotics (Sigma-Aldrich) were dipped into AABC solutions at dilutions ranging from 1:1 (undiluted) to 1:32. The disks were air-dried in a sterile environment. Bacterial suspensions with a turbidity of 0.5 McFarland were prepared, and a lawn culture was inoculated on Columbia plates. AABC-impregnated disks were placed on the surfaces of the inoculated agar plates and incubated for 24 h at 37 °C. The diameters of the inhibition zones were then measured and compared.

The tests were performed in triplicate in three separate experiments.

### Effect of AABCs on microbial biofilms

Bacterial and fungal 24 h and 48 h biofilms were prepared in 96-well flat-bottom microtiter plates, as previously described (79). Following a 24 or 48 h incubation at 37 °C (for bacterial cultures) and 30 °C (for fungal cultures), biofilm samples were washed three times with PBS to remove non-adherent bacteria, after which 200 µL of CA-MHB broth with 2-fold dilutions of AABCs were added. The plates were incubated at 37 °C for 24 h, the well contents were removed via aspiration, and each well was washed three times with PBS.

To determine the OD, subsequently, 100 μL of 0.1% crystal violet solution (Innovating Science, Aldon Corporation, Avon, USA) was added to the wells containing dried biofilms. After 15 min of exposure, excess crystal violet was removed, the wells were washed three times with sterile water, and 150 μL of 95% ethanol was added. The OD was measured at 570 nm (Promega, GloMax reader).

To evaluate the microbial viability following AABC exposure, biofilms (not stained with crystal violet) were scraped thoroughly, with particular attention paid to the well edges (79). The well contents were aspirated, and the total CFUs were determined via serial dilution and plating bacteria and fungi on Columbia agar or Sabouraud dextrose agar, respectively.

### Tumor cell viability assay

Tumor cell viabilities were assessed by performing MTT and LDH assays. H1299 and T98G cells were seeded in 96-well plates (5 × 10^3^ cells/well) in RPMI-1640 medium supplemented with 10% FBS (Gibco) and 1 × penicillin/streptomycin (Sigma Aldrich). After a 24 h incubation, the growth medium was replaced with RPMI-1640 medium containing two-fold serially diluted AABSs without FBS or antibiotics, and the plates were incubated for another 24 h. The MTT assays and LDH were performed according to the manufacturer’s instructions (both Thermo Fisher, Waltham, MA, USA). The tests were performed in triplicate in three independent experiments.

### Evaluating the thermotolerance and protease resistance of AABCs

AABC in microcentrifuge tubes was heated to 70 or 90 °C in a dry bath for 5, 15, or 30 min. The effects of heating on the MICs of AABCs were assessed by comparison with those of the non-heated AABCs.

To evaluate the resistance of AABCs to proteolytic enzymes, AABCs were treated with trypsin (20 µg/mL) or protease K (20 µg/mL) (both from Sigma-Aldrich) for 0.5, 2, 15, or 30 min. Proteolytic enzymes were inactivated using 5 mM phenylmethylsulfonyl fluoride (Sigma-Aldrich), as described previously. The tests were performed in triplicate in three independent experiments.

### Microscopy

The cells were imaged using an Axiovert 40C microscope (Carl Zeiss, Germany) at 32× magnification.

### Protein identification and label-free quantification using ultra-high performance LC (UPLC) coupled with tandem MS

Two aliquots of each culture medium sample (containing 50 µg of protein) were removed for bottom-up proteomics analysis. One aliquot was subjected to standard in-solution digestion procedures, including a 1:1 dilution (v: v) with 0.1 M ammonium bicarbonate in water (final pH of 8). Cys residues were reduced and alkylated using 5 mM DTT and 10 mM iodoacetamide for 1.5 h at 22°C room temperature and 45 min at room temperature in the dark. Sequencing-grade modified trypsin (Promega) was added at a 1:20 dilution (enzyme: protein), and digested proceeded for 16 h at 37 °C on a thermal mixer. Trypsin digestion was quenched with formic acid to a final pH of 2.5.

The second culture medium sample was subjected to full denaturation and high-temperature digestion by adding urea (8 M final concentration). The Cys residues in the urea-denatured samples were reduced and alkylated with 5 mM DTT or 10 mM iodoacetamide at room temperature for 1.5 h or 45 min in the dark, respectively. Endoproteinase LysC/trypsin mixture (Promega) was added to the samples at a 1:20 (enzyme: protein) ratio for 16 h at 37 °C on a thermal mixer for LysC-specific proteolysis. On the next day, the samples were diluted to contain 1 M urea using 0.1 M ammonium bicarbonate to activate the trypsin, after which they were digested for 8 h at 50 °C. The dual-enzyme digestion reactions were cooled to room temperature and quenched using formic acid (final pH: 2.5). Digested and acidified peptides from both digestion protocols were desalted using Pierce Peptide Desalting Spin Columns, dried down, resuspended in 0.1% formic acid in water, quantified by measuring their OD_280_ values using a Nanodrop spectrophotometer (Thermo Scientific), and frozen at −20 °C before analysis.

Desalted peptides were subjected to UPLC-MS/MS analysis using a Dionex Ultimate 3000 RSLCnano UPLC instrument coupled to an Orbitrap Eclipse Tribrid MS instrument (Thermo Fisher Scientific). The peptides were loaded onto a nanoEase m/z Peptide BEH C18 analytical column (Waters Corporation, Milford, MA) 75 µm × 25 cm and separated using a 60 min linear reversed-phase UPLC gradient (solvent A: 0.1% formic acid in water, solvent B: 0.1% formic acid in acetonitrile). The peptides were gradient-eluted directly into the Orbitrap Eclipse using positive-mode electrospray ionization and analyzed using the data-dependent acquisition method for MS/MS acquisition, incorporating higher energy collisional dissociation. Both MS and MS/MS spectra were acquired at high resolution with the Orbitrap instrument.

Peptide identification, protein identification, and label-free quantification were performed using MaxQuant (v2.0.2.0) and an embedded Andromeda search engine. Raw data were searched against the UniProt *Homo sapiens* reference proteome (identifier UP000005640, accessed 02/26/2024) using tryp/P enzyme specificity, a minimum peptide length of 5 amino acids, and variable modifications: oxidation of Met, protein N-terminal acetylation, deamidation of Asn and Gln residues, and peptide N-terminal Gln to pyroGlu conversion. Cys carbamidomethylation was set as a fixed modification. Protein and peptide spectrum match false-discovery rates were set to 1%. The MaxQuant results were uploaded to Scaffold 5 (Proteome Software) for further data visualization and analysis.

### Metabolite identification and label-free quantification using UPLC-MS/MS

We extracted 300 µL culture-medium samples for metabolomics analysis. The samples were subjected to a protein-precipitation step by adding 1200 µL of ice-cold methanol, vortexing the samples for a few seconds, and incubating them at −20 °C for 1 h. The samples were then centrifuged at 16,000 × *g* for 15 min at 4 °C. The supernatants were collected into a pre-weighed 15 mL plastic tube (Corning), mixed with three volumes of Fisher Optima LC-MS grade water, and frozen at −80 °C. The frozen samples were dried using a lyophilizer (Labconco, Shell Freezer, USA) and weighed to determine the dry masses of the extracts. The dried extracts were reconstituted to 10 mg/mL in LC-MS Optima grade water containing 0.1% formic acid and transferred to LC-MS vials before analysis.

The samples were analyzed using a Xevo G2-XS QToF UPLC-MS/MS system (Waters Corporation) in both positive- and negative-electrospray ionization modes. Chromatographic separations were performed on an Acquity UPLC HSS T3 column (150 × 2.1 mm, 1.8 µm particle size, Waters Corporation) with a column temperature of 40 °C. Water and acetonitrile containing 0.1% formic acid were used as solvents A and B, respectively, for positive-mode analysis, and water and acetonitrile containing 10 mM triethylammonium acetate were used as solvents C and D, respectively, for negative-mode analysis. The injection volume was 10 µL. The mobile phase gradient with a flow rate of 0.3 mL/min was maintained as follows: 0–8.5 min: 5% B/D; 8.5–9 min: 98% B/D; and 9.25–12.5 min: 5% B/D. Data were acquired using MassLynx software (version 4.2). Data acquisition was performed in the sensitivity mode using MSe data-independent acquisition for MS2 acquisition. The mass range for MS and MS2 acquisition was 50– 2000 Da. A collision energy ramp of 4–40 V was applied for MS2 acquisition. Leucine enkephalin (400 pg/µL) was used for lockspray mass corrections.

The raw data files from the LC-MS/MS analysis were imported into Progenesis QI software (v. 4.1, Nonlinear Dynamics) for further data processing. The METLIN MS/MS database (v. 1.0.6499.51447) was used for compound identification. The metabolites in the samples were matched against compounds in the database with a 10 parts-per-million (ppm) precursor ion m/z tolerance and a 20 ppm fragment ion m/z tolerance. Compounds were measured and identified for further analysis.

### Statistical analysis

At least three biological replicates were analyzed for each experimental condition in three independent experiments, using samples from at least five donors unless otherwise stated. Data points are presented as the mean ± standard deviation (SD). Bacterial and fungal quantification data were log-transformed (log_10_) before analysis. To avoid taking logs of 0, we added a constant pseudo-count of 0.1, ensuring that all peptides and metabolites were uniquely identified (80).

The statistical tests used are specified in the corresponding figure legends. GraphPad Prism version 10 (GraphPad Software, San Diego, CA, USA) or Excel 10 (Microsoft, Redmond, WA, USA) were used for statistical analyses and illustrations.

## Data availability

If data are to be shared upon request, then the individual along with their contact information must be indicated

## Supporting information

This article contains supporting information.

## Acknowledgements

We would like to thank the Genome Technology Center (GTC) and the Applied Bioinformatics Laboratories (ABL) for providing support and helping with the analysis and interpretation of the data. GTC and ABL are shared resources partially supported by the Cancer Center Support Grant P30CA016087 at the Laura and Isaac Perlmutter Cancer Center. This work has used computing resources at the NYU School of Medicine High Performance Computing (HPC) Facility.

The authors would like to acknowledge the NIH S10 High-End Instrumentation Award 1S10-OD028445-01A1, which supported this work by providing funds to acquire the Orbitrap Eclipse Tribrid mass spectrometer housed in the University of Connecticut Proteomics & Metabolomics Facility.

We would also like to thank Natalija Budimir for working on the manuscript.

**Funding and additional information**

**Conflict of interest**

**References**

## Abbreviations

AABC: anticancer and antimicrobial bioactive compound
ANOVA: analysis of variance
ATCC: American Type Culture Collection
CFU: colony-forming unit
CLSI: Clinical and Laboratory Standards Institute
FBS: fetal bovine serum
CA-MHB: Cation adjusted Mueller Hinton Broth
LDH: lactate dehydrogenase
MFC: minimum fungicidal concentration
MIC: minimal inhibitory concentration
MTT: 3-(4,5-dimethylthiazol-2-yl)-2,5-diphenyltetrazolium bromide
NSCLC: non-small cell lung cancer
SD: standard deviation
SEM: standard error of the mean
TezR: Teazeled receptor
UPLC: using ultra-high performance liquid chromatography

---

**Supplementary Figure 1. Antimicrobial activity of AABCs derived from Leukocyte-Tells.** (A) Inhibition of 48 h old bacterial biofilms with undiluted supernatants containing AABCs derived from Leukocyte-Control and Leukocyte-Tell cultures. The combined results of three independent experiments are depicted. The bar graphs represent the mean ± SEM. Statistical significance was determined via two-way ANOVA. (B) Decreased CFUs of 48 h old fungal biofilms treated with supernatants containing AABCs derived from Leukocyte-Controls and Leukocyte-Tells. The combined results of three independent experiments are depicted. The bar graphs represent the mean ± SEM. Statistical significance was determined via two-way ANOVA.

**Table S1. Differentially produced metabolites identified in the supernatants of Leukocyte-Tells.**

**Table S2. Non-peptide metabolites uniquely found in the supernatants of Leukocyte-Tells**

