## Supplementary figures and images for "The Universal Receptive System acts as a novel regulator in the production of antimicrobial and anticancer bioactive compounds by white blood cells"

### Supplementary Figure 1

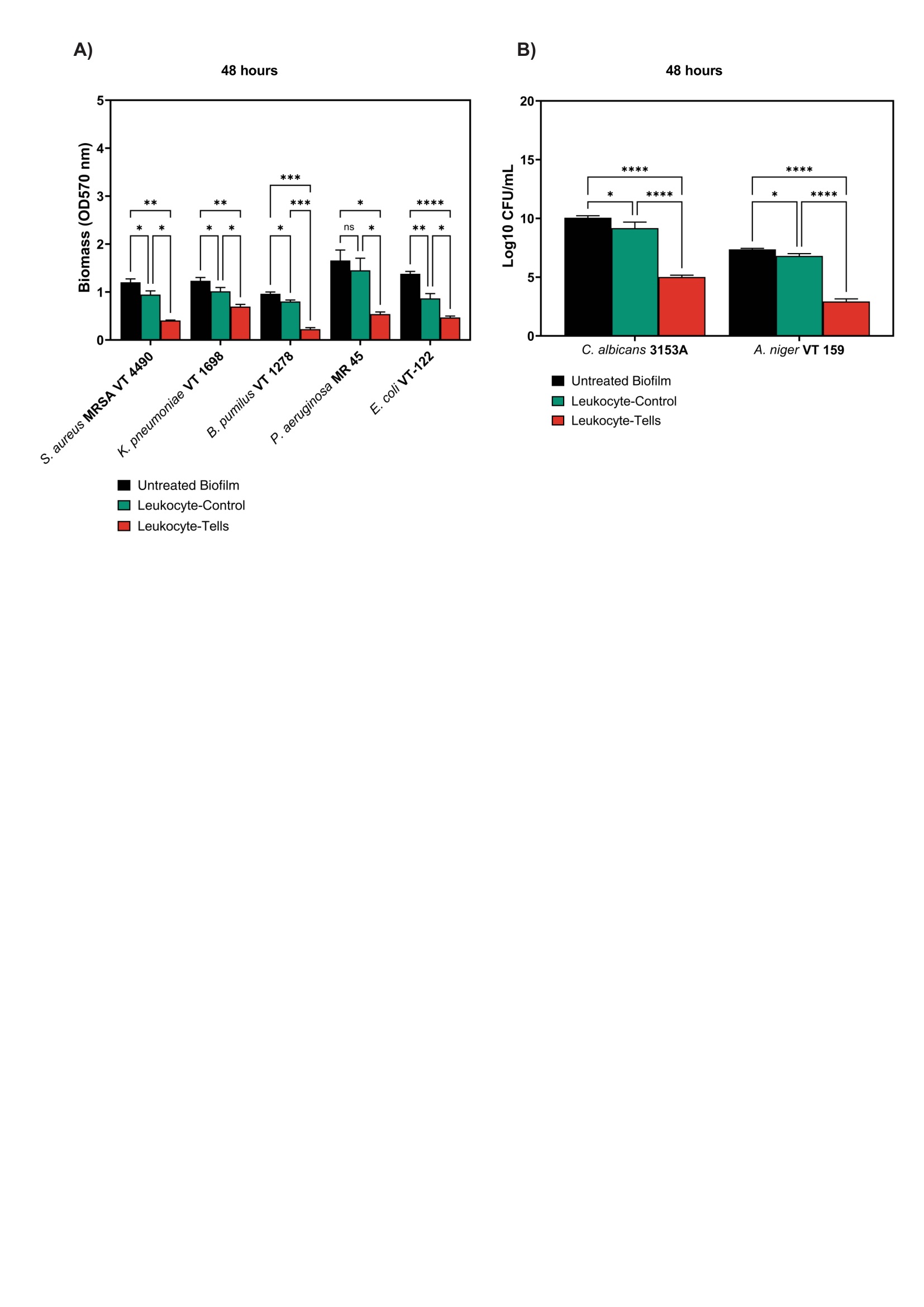
